# The fractured landscape of RNA-seq alignment: The default in our STARs

**DOI:** 10.1101/220681

**Authors:** Sara Ballouz, Alexander Dobin, Thomas Gingeras, Jesse Gillis

## Abstract

Many tools are available for RNA-seq alignment and expression quantification, with comparative value being hard to establish. Benchmarking assessments often highlight methods’ good performance, but are focused on either model data or fail to explain variation in performance. This leaves us to ask, what is the most meaningful way to assess different alignment choices? And importantly, where is there room for progress? In this work, we explore the answers to these two questions by performing an exhaustive assessment of the STAR aligner. We assess STAR’s performance across a range of alignment parameters using common metrics, and then on biologically focused tasks. We find technical metrics such as fraction mapping or expression profile correlation to be uninformative, capturing properties unlikely to have any role in biological discovery. Surprisingly, we find that changes in alignment parameters within a wide range have little impact on both technical and biological performance. Yet, when performance finally does break, it happens in difficult regions, such as X-Y paralogs and MHC genes. We believe improved reporting by developers will help establish where results are likely to be robust or fragile, providing a better baseline to establish where methodological progress can still occur.

## INTRODUCTION

A major computational challenge in RNA-sequencing is the quantification of gene expression. At present, the most commonly used approach consists of mapping RNA-seq reads to a reference, followed by calculation of transcript or gene expression levels (see comprehensive list here (1)). Some newer methods skip a definitive mapping step to generate counts that pass through the probabilistic nature of assigning a given read to any single genomic location (i.e., quasi-mapping (2), pseudo-alignment (3)). Short read sequences, splicing, as well as the abundance of paralogous sequences in the genome make definitive alignment challenging. Furthermore, the pervasiveness of overlapping isoforms (owing to alternative splicing as well as alternative transcription start/termination) make the quantification non-trivial with most mapping tools relying on sophisticated statistical models to estimate the likelihoods of reads originating from various transcripts. Notwithstanding similarity in output of these tools - or because of it – debate as to best practices remains vigorous. Ideally, this would mean that the field as a whole has converged on a reasonable, robust, and biologically informative set of practices to at least characterize gene expression. Unfortunately, what is biologically informative is very hard to assess in a general way. Understandably, this has led to a focus in assessment on comparatively straightforward metrics or simple biological tasks, accentuating positive results but potentially reflecting overfitting in assessment if intended to generalize to novel data or new experimental systems.

As novel tools are developed, neutral or independent evaluations compare them to currently available methods, typically done as a benchmarking exercise. Generally, a benchmark will be designed to critically evaluate tools based on a number of criteria such as accuracy, reproducibility, and efficiency. Ideally, it will pinpoint the strengths and weaknesses of the tool which are likely to be relevant to its user base, allowing them to make informed decisions dependent on their specific data or experimental design (4). Although not entirely neutral, as much as that is the intent (5), the goal is to be fair and comprehensive in assessment. Yet, purely technical metrics like mapping efficiency and rates do not trivially relate to any specific biology under investigation. Alternatively, biologically relevant metrics, such as known differential expression under some change in condition, will tend to involve simple experimental systems (in order to have a clear answer) and may not generalize to more complex systems (6,7). To some extent these issues are visible in most outcomes of benchmarking endeavours and are therefore appreciated by the field. Some call for exhaustive validation of all novel tools in the hope that this will give an equal ground for the comparison (8), while others argue for more representative (e.g., across multiple types of tests) and objective (e.g., blinded or double-blinded) comparisons to work as guides for choice (9).

Here, we perform an assessment of assessments, focusing on the characterization of gene expression from RNA-seq data. We observe that most alignment tools work fairly well, but can fail (comparatively) under certain parameters and on different datasets. In order to explore these limits, we perform a targeted assessment on a single tool, STAR (10), and one well-characterized dataset (GEUVADIS (11)), and exhaustively evaluate its performance across its parameter space, providing an in-depth view of one algorithm’s response to challenges. In parallel to these results, we assess a wide corpus of data across common pipelines to validate the presence of the same phenomena revealed by probing STAR’s performance. We first use commonly reported metrics in assessments and then more biologically oriented tests to score the performance of the alignment on both per sample and per experiment basis. We find that changes in alignment parameters within a wide range have very little impact even technically, which in turn has very little impact on biology. However, dramatic failures eventually occur and we conclude that it is likely that the real performance of algorithms is their utility across these hard-to-generalize cases. We also find that long-standing limitations in functional annotation among genes impose a major limit on assessing the utility of methods.

## MATERIAL AND METHODS

### Benchmarking studies summary

Data was parsed from the omicstool (1) website [https://omictools.com/, accessed November 2016], a curated collection of bioinformatics tools. To pick benchmarking papers, we used those listed as benchmarking papers on the omicstools site [date November 2016] and papers from RNA-seq blog [http://www.rna-seqblog.com/, accessed November 2016]. Mapping rates and correlations were extracted from the papers supplementary tables, if available. A total of 11 papers had aligner mapping rates, and a total of 5 papers had quantifier correlation results. These were manually curated and can be found in the supplement (**Table S1**).

### Datasets

We perform all our assessments on publically available RNA-seq data. We used the GEUVADIS (11) dataset, which consists of 462 lymphoblastoid cell line samples (LCLs), split between 246 female, and 216 male. The GEUVADIS data set is available at the European Nucleotide Archive (accession no. ERP001942). For our meta-assessment of RNA-seq expression databases, we downloaded data from three databases. First, Gemma (all datasets available as of Nov 2016), a quality controlled database that used bowtie2 (12) for alignment and quantified with RSEM (13). A second database was ARCHS^4^ (BioRxiv: https://www.biorxiv.org/content/early/2017/09/15/189092) (v1 downloaded Jan 2018), that uses kallisto (3). A final database was recount2 (v1 downloaded Jan 2018) (14) that used the Rail-RNA pipeline (15). We selected a subset of 57 experiments (**Table S2**) totalling 3,405 samples that were common among the three and that had at least 20 samples per experiment. We used MetaSRA (16) to obtain sample labels and metadata through their SQLite database [http://metasra.biostat.wisc.edu/download.html].

### Alignment parameters

STAR version 2.4.2a was run on the FASTQ files of the GEUVADIS datasets. We used genome version GRCh38.p2 and GENCODE version 22 (17). The parameters changed were the minimum alignment score (minAS, parameter: --outFilterScoreMinOverLread) and number of mismatches (numMM, -- outFilterMismatchNmax). The minimum alignment score was varied to range between 0.55 and 0.99. The number of mismatches allowed was varied to range between 0 and 9. We downsampled reads with a local script by sampling across the mapped reads at random. RSEM (version 1.2.28 (13)) was run to quantify the expression levels of the GEUVADIS dataset. We considered both FPKM and TPM. Links to scripts can be found in the supplement.

### Alignment metrics and comparisons

To compare outputs of the alignments, we used the same metrics as reported most commonly by the benchmarks: read mapping rate, and sample-sample correlations using Spearman’s rank correlation coefficient. We report uniquely mapped reads to features as our mapping metric, which was taken from the STAR “ReadsPerGene.out.tab” file. Counts for each gene were collected from the fourth column of the “ReadsPerGene.out.tab” file from STAR. Total mapped counts to features were calculated as the sum of all gene counts. Total input reads were taken from the “Log.final.out” file from STAR. We calculate sample-sample correlations with Spearman’s rank coefficient using the counts per million (CPM) for each sample. This normalizes the data within a sample, and allows comparison across samples with different sequencing depths. The CPM of gene *i* (*CPM_i_*) is the count of gene *i* (*c_i_*) divided by the total counts 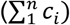 multiplied by 10^6^:

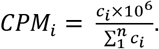

For the database comparative assessment portion, we calculate fraction mapped for each sample by using the read counts from each database, and estimating the fraction mapped from total input reads. We perform gene level sample-sample correlations (Spearman correlation coefficient) between each samples expression levels across the same subset of 31K genes for each pair of the three databases.

### Differential expression analysis

We test for the effect of the alignment on the ability of the sex-specific samples to determine which sex-specific genes are differentially expressed. To do this, we calculated the log2 fold change for each gene (*FCi*) between female and male samples, as the log2 of the average CPM over female samples divided by the average CPM over male samples. We calculate a p-value using the Wilcoxon rank-sum test (wilcox.test in R) and adjust for multiple hypothesis tests using Benjamini-Hochberg (p.adjust in R). The degree of differential expression for gene *i*, is the average rank it takes over both fold change (*FC_i_*) and the FDR (adjusted p-value, *q_i_*):

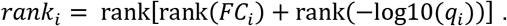

Treating known sex-specific genes as positives, we can calculate an AUROC for the rank these genes take in the sex-based differential expression:

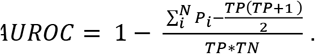

Here, *Pi* is the rank of the positive genes being predicted, *TP* is the number of positives, *TN* is the number of negatives, and *N* the total number of genes. An AUROC of 0.5 is random, 0.7-0.8 is quite good, and anything greater than 0.9 is excellent (within expression data). The test genes sets we use are the Y chromosome genes (our positive test set, 594 genes), sex-specific DE genes (18) (126 genes), and X chromosome genes (negative test set, 2476 genes). We consider genes on the X chromosome as a negative test set since there are regions on both the X and Y that overlap (pseudoautosomal region) and genes that are homologous. As a control experiment, we shuffle the samples (mixing males and females), and repeat our tests.

As a measure of sex specificity, we then calculate the fraction of chromosome Y genes detected *S_j_* in sample *j* as the sum of Y chromosome genes with expression level greater than zero divided by the total number of Y chromosome genes. Fractions closer to 0 are more female, and those closer to 1 are more male.

### Biological replicability analysis through co-expression

Co-expression is a measure of gene-gene co-variation that has biological implications including co-regulation and co-functionality. In previous work (19), we used this concept to detect how well an experiment replicates by measuring how well it has retained well known co-expression patterns. Thus, to measure the biological replicability of an experiment, we ran AuPairWise (19) using default parameters (25% noise, 100 repeats per sample) to calculate co-expression scores. From the two AUROCs that are given as output, one for the stoichiometric pairs and one for the random pairs, we calculate a co-expression score for each sample as the log ratio:

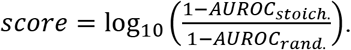

This score measures the degree to which previously observed co-expression relationships are present in the dataset relative to random co-expression. Since we use a set of primarily protein complexes which are highly co-expressed in all other data, this assessment is general biologically. Each AUROC measures how well a sample which whose expression values have been perturbed can be identified based on its disrupted co-expression (i.e., which samples are outliers when two genes which are generally correlated are plotted against one another).

## RESULTS

### Alignment benchmarks on average show similar performances

Transcriptome analysis through RNA-seq requires expression levels to be quantified and summarized on a per gene (or isoform) basis, with many pipelines available to accomplish this task (summarized in **Figure 1A**). First, a sample is processed and sequenced, returning millions of reads/fragments. This is then followed by the mapping stage - reads are aligned to a reference genome or transcriptome - and then quantified to measure expression levels in genes or transcripts. Each step has its own source of errors, which affect the difficulty of measuring the transcriptome accurately. Once mapped and quantified, the transcriptome is used for a multitude of purposes: detecting whether a gene is expressed, differential expression between conditions or co-expression across conditions (**Figure 1B**). We conducted a survey of all tools published between the advent of RNA-seq in 2008 until 2016, and find a steady increase over time (**Figure 1C**), with multiple new tools published yearly. Each new method developed motivates a benchmark assessment between the newly developed tool and, typically, the most popular methods (between 5 and 10 tools), yielding recurrent assessment of a subset of tools. In these assessments, the choices of performance metric to evaluate the outcome vary, and we found the two most common metrics to be sample specific, and were the number of unique reads mapped and correlations of expression profile with either a reference standard (ideally a gold standard) or among the competing tools. We will refer to these metrics as the “fraction mapped” and the “correlation”.

**Figure 1.**
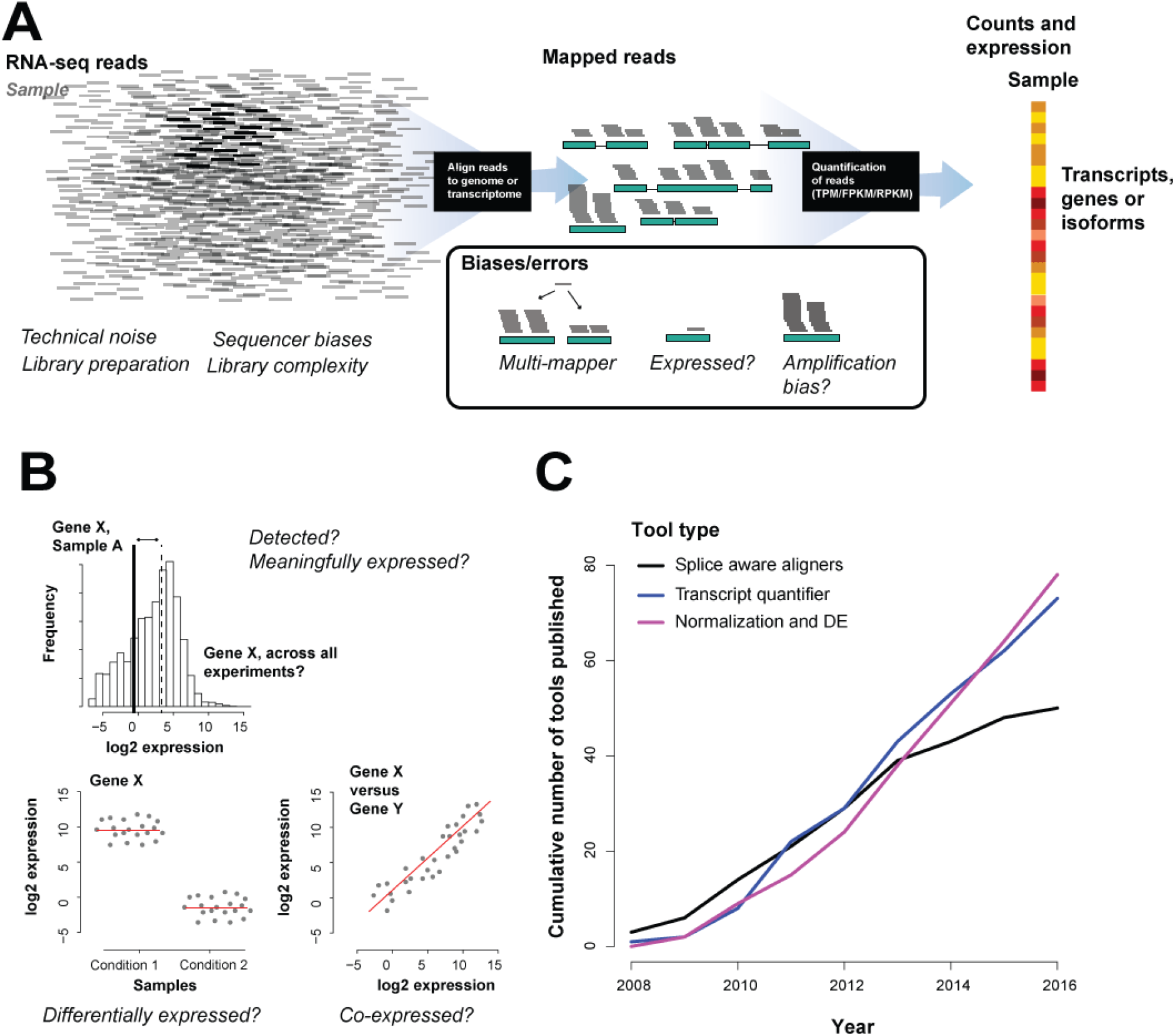
Summarizing RNA-seq alignment tools. (A) RNA-seq alignment: typical experiment + sources of error. (B) Transcriptomics asks three broad questions which include transcript detection, differential gene expression and gene co-expression. (C) Growth and performance of RNA-seq tools - cumulative number of tools shows steady growth across all tool types.

We compared the outcomes based on those two metrics from published assessments within the last few years, with the aim of determining their consistency. To provide a broad overview of comparative performances, we calculated the reported fraction mapped performance for each tool, ordered by the number of tests (**Figure 2A**). The performances of a tool appeared to be variable across studies, as shown by the wide range of values for each tool. Summarizing this effect by calculating variance in fraction reads mapped across studies for each method gives a mean value of ~0.15 (SD) but also shows approximately two modes. The higher mode tends to be occupied by the most tested methods, and consists of cases where they do exhibit a high degree of variability in performance (higher mode in purple violin plot, SD+/-~0.35 **Figure 2B**). Methods with lower variances (SD+/-~0.04) were tested fewer times; both the mean performance and its variance were correlated with the number of tests (rs=-0.5 and rs=0.55, respectively). If we instead assess the variability in performance within each assessment (across tools), we obtain a similar total variability (~0.15 SD) and another very heavy tail, reflecting an apparent bimodal split (SD+/-~0.05 and ~0.43), suggesting there are once again two pervasive classes of result in the assessments. Firstly, there are a set of tests where all methods assessed have fairly similar performances on the same dataset and this is indicated by the low SDs in the violin plots (53% with mean SD+/-~0.15, by dataset). The second type of result is where the methods perform quite variably on a dataset, implying that some of the methods may be optimized (relative to other methods) for a particular dataset.

**Figure 2.**
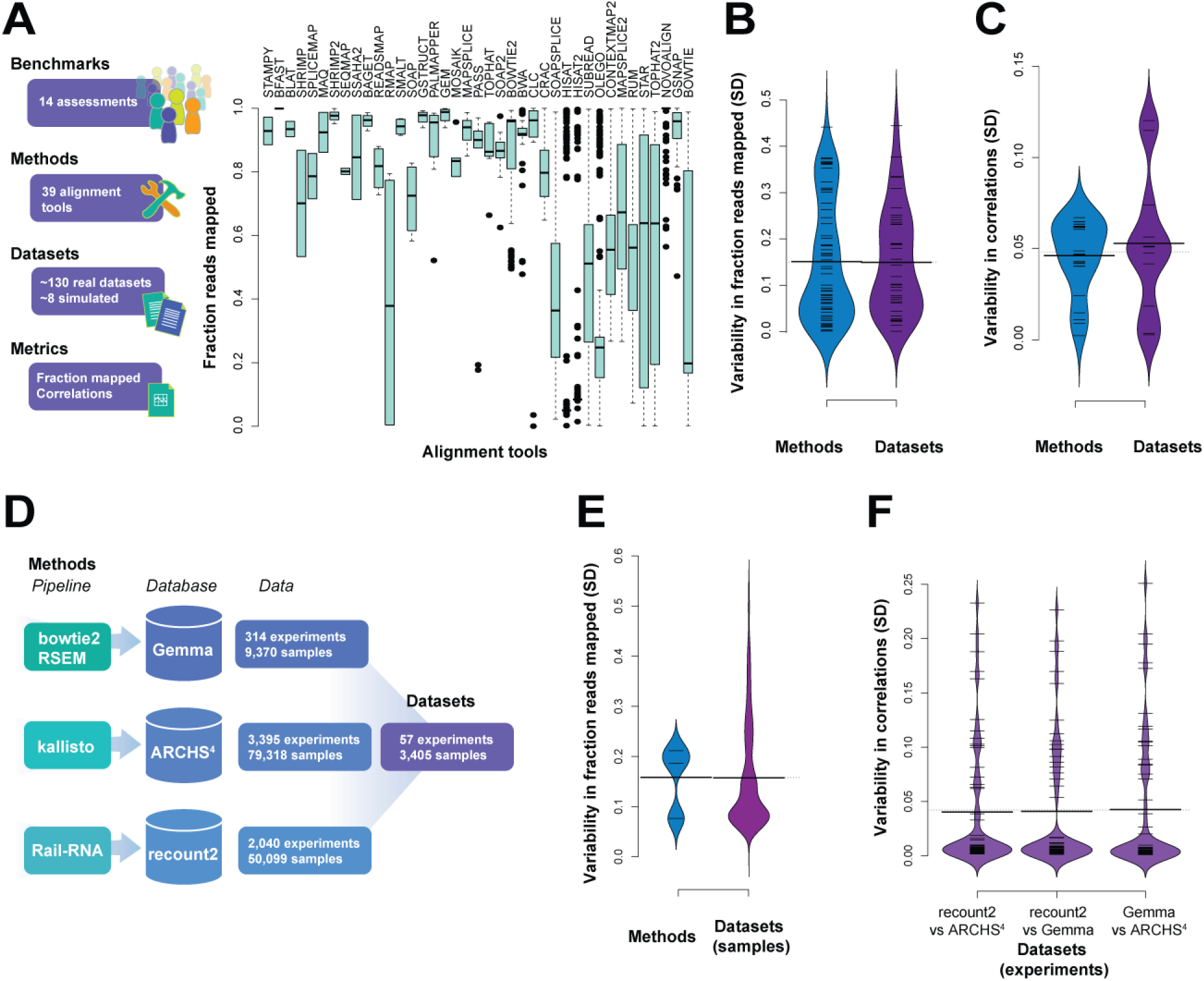
Summary of previous studies of RNA-seq alignment evaluations. (A) Fraction of reads aligned for aligners across different tools, datasets (real and simulated) and parameters, ordered by number of times assessed. (B) SDs of the fraction mapped comparing averaging by dataset or by method. (C) Correlations of the relative expression levels of the output of different quantifiers in a subset of 5 benchmarks. (D) Summary of databases and pipelines used in the global assessment. Overall, 57 experiments and 3,405 samples were used in the assessment. (E) SD of fractions reads mapped per database and those across the three databases (per sample) (F) and correlations of the expression levels between the different databases (SDs per experiment).

Beyond even the evident variation in performance, the very definition of the “fraction mapped” metric may vary significantly between the studies. This metric is affected, for instance, by discordantly mapping read pairs, clipping of the reads, maximum number of allowed mismatches, etc. Since the benchmarking is usually performed with default aligner parameters, the aligner filtering preferences may collide or harmonize with the metric choices, thus creating assessment biases even within one study. The “fraction mapped” metric is also easy to game provided we permit a greater number of mismatches. However, notwithstanding this potential for metric misrepresentation, we obtain a quite similar narrative when looking at benchmark outcomes using correlations as the reported metric. While we do see tools tend to perform more similarly within each tested dataset (SD+/-~0.05, grey violin plot in **Figure 2C**), performance of a given method across datasets remains highly variable with a similar bimodal split to previously (purple violin plot **Figure 2C**).

In order to characterize the field-wide process of RNA-seq alignment more comprehensively, we looked to three largescale databases with mapped RNA-seq data across thousands of datasets (**Figure 2D**). This provides us with a view of the global data landscape, and also one over which we can still explore method variability by exploiting overlaps between the databases (each of which use distinct pipelines, see methods). We treat each overlapping experiment (across its samples) present in all three databases as a distinct evaluation. Variability in fraction reads mapped is very similar to the benchmark evaluation across both methods (SD+/-~0.16) and datasets (SD+/-~0.16) within this broader survey (**Figure 2E, Figure S1**). There is a very modest attenuation of the variability in fraction reads mapped across data. Interestingly, the variability in correlation has a similar range of values to the benchmark data not just in aggregate (SD+/-~0.04 for each comparison on a per sample basis, but when averaged over each experimental dataset, **Figure 2F**). This is consistent with the view that variation in biological context – likely to be held constant within an experiment but varying between - substantially affects the apparent efficacy of RNA-seq alignment.

### The limits of benchmarking metrics

Our summary of benchmarks suggests that tools perform too variably to generalize their performance ability across datasets: a tool can be optimized to perform well on a particular dataset but has breakpoints – parameters or data that cause the software to perform poorly. To characterize these breakpoints and quantify the effects of these in a benchmark, we perform a controlled assessment using STAR (10) on a single dataset GEUVADIS (20), permitting a more exhaustive evaluation of performance dependency than is typically feasible. To estimate the performance space over crucial parameters, we focused on the --outFilterScoreMinOverLread parameter in STAR, which controls the minimum alignment score normalized to the read length, which we denote as minAS. There is no clear a prior expectation about how reducing the minimum permitted quality of mapped reads will affect either technical or biological efficacy – more data often compensates for less stringent filtering.

Looking once again to the first of our benchmarking statistics (“fraction mapped”), we observe a range of mapping values for each sample for the given alignment score parameter (width of plots in **Figure 3A**), and fairly stable average mapping for the experiment (average of violin plot **Figure 3A**). As expected, the less stringent this score, the higher the fraction mapped metric (e.g., minAS=0.55 has a fraction mapped of 74% SD+/-~0.04). The lowest boundary was selected to allow only concordantly paired alignments. If the minAS is ≤0.5, the single-end and discordantly paired alignments will be returned as output, which significantly increases the percentage of mapped reads, and at the same time increases the rate of mis-alignments. We also see that the fraction mapped is similar for most of the parameters, but then drops considerably near the very conservative scores (anything greater than >0.90), with only 44% (SD+/-~0.07) of the reads mapping at the far end (minAS=0.99), where only alignments almost perfectly matched to the reference are allowed. This decrease in performance was unequally distributed across the samples in the data with some samples dramatically affected (e.g., ERR188390 75% to 17% SD+/-~0.21) and other much less so (e.g., sample ERR188428 43% to 25%, SD+/-~0.06). This parallels our observation in the benchmarking summary of occasional breakpoints not being evenly distributed across data (or methods). Although they are comparatively easy to detect in this focused evaluation, they may well drive variability in performance where defaults are used on data which happens, idiosyncratically, to render those defaults less useful for particular tools.

**Figure 3.**
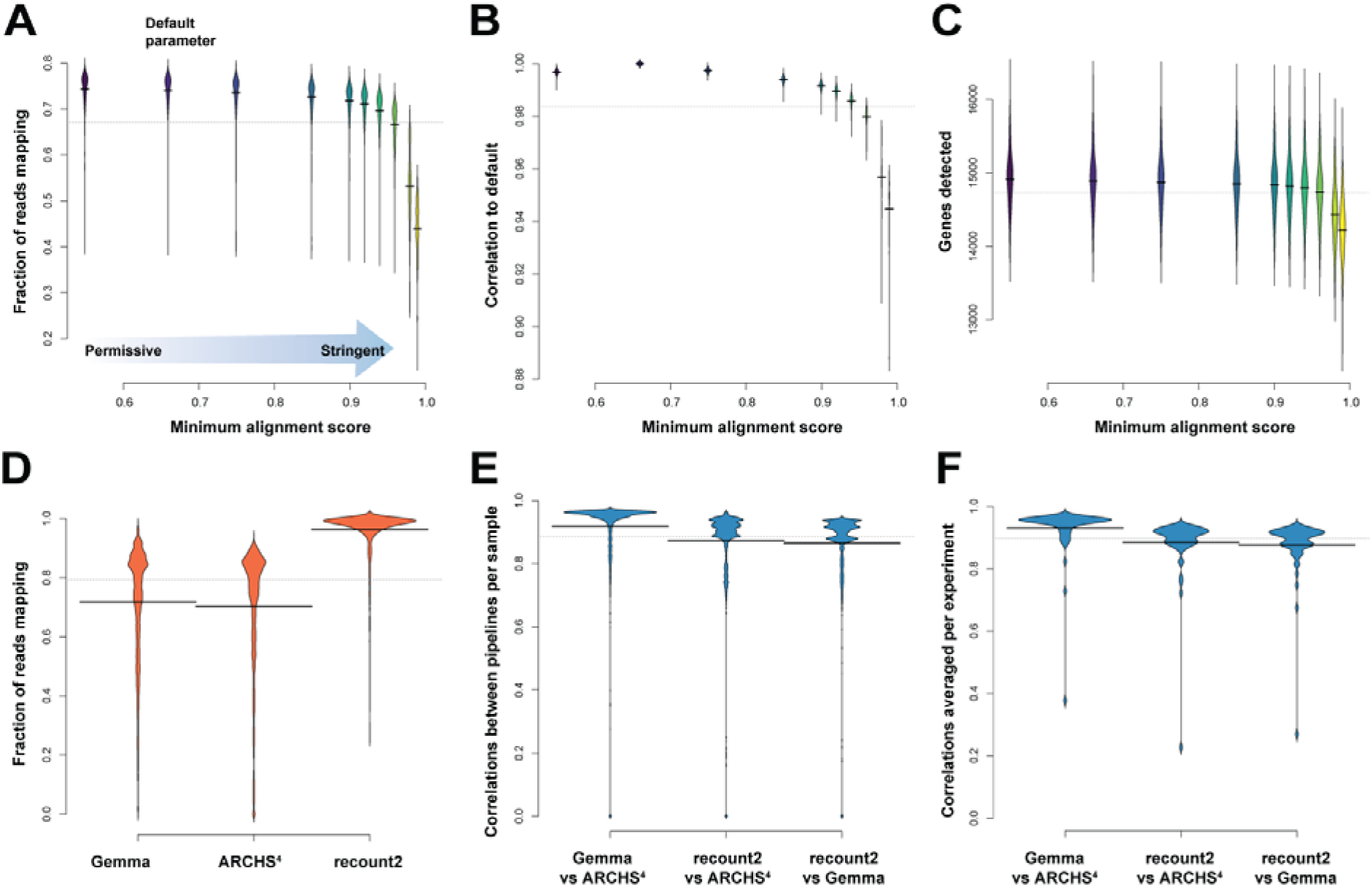
Comparative performances across metrics. (A) Mapping rates for 462 samples from the GEUVADIS dataset varying a single parameter, minAS (--outFilterScoreMinOverLread). The default is 0.66. The most permissive parameter we tested was 0.55, and the most stringent at 0.99. (B) Spearman correlations between a sample and its default counts across the same parameter space. (C) Number of protein coding genes detected for each alignment parameter. (D) In parallel to (A), the distribution of fraction mapped metrics for 3,405 samples in the Gemma, ARCHS^4^ and recount2 expression databases. (E) In parallel to (B), the distributions of the correlations between genes for each sample (pairwise across databases) and then (F) averaged per experiment. Recount2 has quite higher mapping rates, but the samples correlate less with the other two databases, as shown in (E) per sample and (F) averaged per experiment.

For our second benchmarking statistic, we took the default alignment parameter score (minAS=0.66) as our reference against which to calculate correlations. For each sample, we calculate the Spearman correlation between gene expression calculated with minAS=0.66 and gene expression calculated with other minAS values. The correlations are high (0.88<rs<0.99, **Figure 3B**). However, like with the previous metric, the stricter our alignment parameter, i.e., the more conservative we are, the lower the correlation. Thus, in this case, better quality does not compensate for smaller quantity. This is also clearly evident in another commonplace characterization of performance, “gene detection”, defined as the number of genes overlapped by at least one uniquely mapped read. In our assessment here, gene detection also varies between the different parameters, ranging between 12.5K and 16K genes (**Figure 3C**). Similar to the previous two metrics, the detection is similar across the parameters until the most stringent parameter choice. Looking at how these performance metrics compare to those from the three RNA-seq databases, we find that STAR at default on the GEUVADIS dataset performs well within the average range for both the fraction mapped (**Figure 3D,** mean fraction mapped 70%-96%) and correlation metrics assessed (**Figure 3E-F,** mean correlations 0.86-0.93). Also, the worst performing parameters of STAR fit within these distributions, as there exist samples that do fail under the pipeline defaults (e.g., mapping at less than <1%).

### Sex-specific gene expression as a metric of alignment quality

In the preceding analyses, the effect of alignment quality can be trivially measured and understood, however, these analyses speak to the technical rather than biological aspects of the experiment. One of the simplest versions of assessing the latter involves using sex as a differential condition (7). In this case it is easy to assess at least a gold standard set of false positives and false negatives, since the genes differentially expressed between males and females are predominantly genes on the Y chromosome (all or nothing), and known sex specific transcripts such as *XIST*. To evaluate the impact of parameter changes on differential expression, for each value of minAS, we selected the top differentially expressed genes (DEGs) based on |log2 FC| > 2 and FDR <0.05 and calculated the concordance with the default minAS results. In the case of the minAS=0.99 parameter (**Figure 4A**), we observe 45 genes in common out of 58 detected under the default parameters (hypergeometric test p~0). As we increase the mapping stringency (starting from our minimum value 0.55), fewer genes are detected as DE, and the overlap decreases too, falling from a maximum of 97% down to 77% (**Figure 4B**). To be more specific to the task of selecting the correct sex-specific genes, we can calculate an AUROC for a variety of test sets out of the entire gene list (ranked by degree of DE). Our expected DEG sets include (i) positive DEG set of the Y-chromosome genes; (ii) positive DEG set of known sex DE genes (18); (iii) and a negative DEG set, which we chose as all X-chromosome genes, because of the regions homologous with the Y, and hence the potential for technical error driving signal between sexes while still having a strong expectation for largely equivalent dosage (21,22). For the Y-chromosome genes, the AUROC is ~1 (purple line **Figure 4C**); ranking genes by DE yields perfect detection of this set as all Y-chromosome genes are ranked close to first. The discriminatory power of the broader sex DE gene set is weaker, but still fairly high for the positive test (AUROC=0.86, teal line **Figure 4C**). The X-chromosome genes are close to random (AUROC=0.56, green line **Figure 4C**), as expected. If we shuffle the samples, mixing males and females as a negative control task, we find the expected AUROCs now closer to 0.5 for all test gene sets (dashed lines in **Figure 4C**). Repeating this analysis across the other alignment parameters (**Figure 4D**) we find almost indistinguishable AUROC values for all gene sets for both positive and negative controls tasks. Even though the mapping statistics and correlations strongly depend on the minAS parameter (see previous section) the downstream biological application (sex-DE) is not affected significantly by the choice of this parameter.

**Figure 4.**
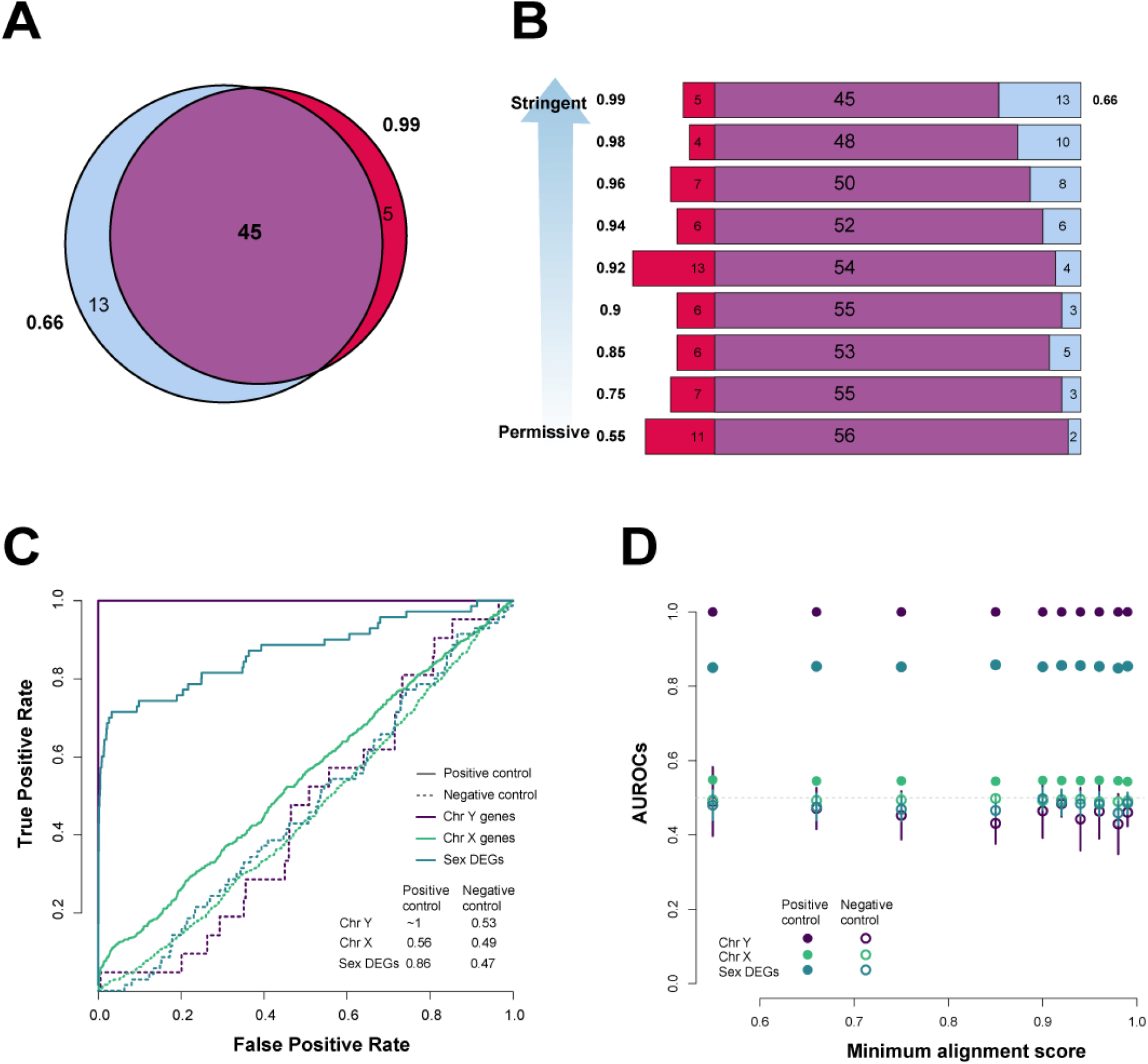
Differential expression by sex in the GEUVADIS dataset. (A) Overlap of all DE genes (FDR <0. 05 and |log 2 FC| >2) for the default minAS= 0.66 and extreme minAS=0.99 (B) and then across parameters with the default parameter. (C) ROCs for each set using average rank of the fold change and the adjusted p-value (see methods). (D) AUROCs across the minimum alignment parameters for the two gene sets tested for both negative and positive controls. These are averaged across 10 runs of the analysis, sampling 200 males and 200 females from the totals.

### Easy-to-validate sex-specific gene expression is too easy to test alignment

To reconcile the minimal change in signal in the DE task with the substantial change in gene detection and expression levels, we characterized the samples on a per gene basis. In this dataset, since half the samples are male, there should be no expression in half the samples of the Y chromosome genes, but we find genes with expression in more than 200 samples, potentially implying wrong mapping/alignment, given proper sample labelling and QC. There are female samples in which some reads map to the Y-chromosome genes (as many as 23 genes, red distributions in **Figure 5A**). Some of these are pseudogenes (e.g., *EIF4A1P2* and *PSMA6P1*), detected in close to all samples. Protein-coding genes detected in over 75% samples share known paralogs on the X chromosome (**Table 1**). For instance, ribosomal protein S4, Y-linked 1 (*RPS4Y1*), is detected in 433 samples and is the only ribosomal protein encoded by more than one gene (ribosomal protein S4, X-linked, *RPS4X*). If we measure the fraction of Y-chromosome genes detected and compare it to the average expression, we find low average expression, with much lower CPM in females than in males (red dots females, blue dots males **Figure 5B**). So although there is incorrect alignment, we still are able to detect male-ness over female-ness in expression and in differential expression, as demonstrated in the previous DE task. On a per sample basis, while there are a few male samples with low average expression, there are almost always enough genes detected in these samples to ‘classify’ them as male. This classification only improves as we increase stringency across the minAS parameter; we see positive correlations between fractions detected and average expression for female samples as we vary minAS, and negative correlations for male samples. We demonstrate this in a female sample (top panel **Figure 5C**) and male sample (bottom panel **Figure 5C**), and all samples (**Figure 5D**). Thus, presence or absence call on a per gene basis is rendered more accurate with increasing stringency but there is little improvement in differential expression on a per sample basis and across samples there is no change in aggregate performance at all.

**Table 1.** Genes on the Y chromosome detected in at least 75% of samples

| Gene symbol | Paralog(s) X | Number of samples |
| --- | --- | --- |
| *EIF4A1P2* |  | 461 |
| *PSMA6P1* |  | 461 |
| *RPS4Y1* | *RPS4X* | 433 |
| *DDX3Y* | *DDX3X* | 419 |
| *VDAC1P6* | *VDAC1P1, VDAC1P2, VDAC1P3* | 383 |
| *RPL26P37* |  | 373 |
| *KDM5D* | *KDM5C* | 368 |
| *EIF1AY* | *EIF1AX* | 365 |
| *TXLNGY* | *TXLNGX* | 364 |
| *USP9Y* | *USP9X* | 351 |

**Figure 5.**
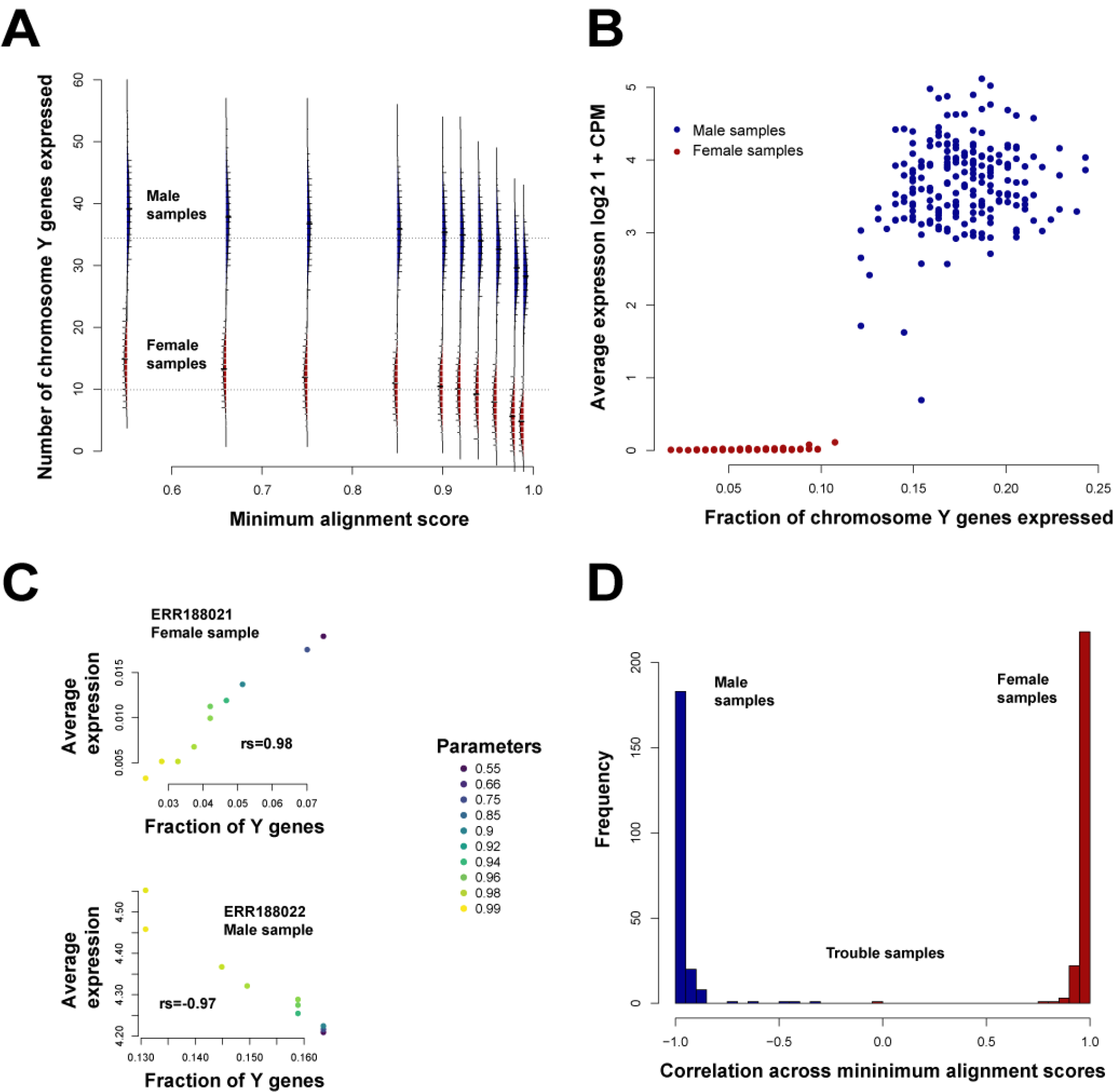
Performance details for sex-specific differential expression. (A) Distribution of number of Y chromosome genes expressed in female (red) and male (blue) samples across all parameters. (B) Average expression of these genes, and a measure of the female-ness and male-ness of the samples (fraction of chromosome Y genes detected) at default minAS=0.66. Samples colored by sex (females in red, males in blue). (C) Two sample experiments showing the change in expression and number of Y genes detected as minAS changes. Top panel is a female sample showing more genes are detected as expressed with higher expression levels as we decrease the minAS (rhos=0.98), but this relationship is inverted for a male sample (bottom panel, rho=-0.97). (D) Plot of the correlation between average expression and Y genes detected across the minAS for all samples showing the same trends as in (C). There are a few samples where these relationships are much weaker.

We then quantified the alignments from the default parameter set and a stricter alignment parameter (minAS=0.99) with RSEM (**Figure S2**). We find that RSEM quantification exacerbated the problem of misalignment, suggesting that error propagates for these edge cases. Even with stricter parameters, once quantified, Y genes with zero counts were suddenly found as expressed (**Figure S2, Figure S3**). Multimapping reads appear to be one of the reasons for this as these reads map to genes that are likelier to be paralogs, homologs, or within gene families. Genes that have multimappers will need to be interpreted with caution, either ranked lower in results space or weighted by the confidence of mapping using fraction of multimappers.

Once more we return to the meta-assessment across expression databases and look at X-Y alignment in the 3,405 samples. Only 761 samples had sex annotated (see methods). As in our detailed assessment, we do find alignment to the Y chromosome genes in the female samples (**Figure S4A**). Indeed, the importance of an appropriate threshold (allowing for alignment error) is highlighted by the fact that the fraction of the Y chromosome expressed at all (non-zero) is quite similar in male and female samples. An appropriate threshold (1-10 CPM) mostly correctly distinguishes meaningful Y chromosome expression (in males) from noise (in females); however this can vary depending on pipeline as shown in two example experiments (e.g., GSE60590, GSE61742; **Figure S4B**). Indeed, comparing the different values databases give to a given sample for fraction of Y chromosome genes expressed shows much greater scatter in female samples than male, as might be expected for low-expression (and errors **Figure S4C-F**). The scatter present even for male samples is potentially surprising, however (**Figure S4E**).

### Co-expression as a genome-wide metric for assessing alignment

Our sex-specific analyses raise the possibility that technical metrics of QC and biological value are, if not in conflict, at least less clearly related than might be hoped. However, because the previous sex-differential task is a comparatively easy one, it is not well suited to detect subtle differences in overall alignment performance, other than the targeted sex-specific genes. One potential solution to this specificity problem is to use a metric capturing very broad-based biological properties present in most expression data whether it is a target of the experimental design or not. Co-expression has historically fit within this category, being both presumed and assessed to be present in essentially any biologically real data (23,24). Co-expression is measured through the correlation (or related) of the expression profiles of a gene pair, and can be measured between all gene pairs within an experiment. As such, we can use co-expression as a replicability score to assess alignment in a very broad and still biologically relevant sense. We calculate a co-expression replicability score using the aggregate signal from gene pairs across the genome that we expect to show high co-expression ((19), see methods). Negative scores indicate observed and specific co-expression (with scores below −1 being excellent), while positive scores indicate the sample exhibits little to no biologically specific co-expression. With the default parameter (minAS=0.66), we observe an average score of −0.52 (SD+/-~0.24), and scores ranging between −0.995 and 0.30. We see again a similar scores with all other parameters (**Figure 6A,** average scores between −0.49 and −0.52). The spread of values for each minAS is broad across samples, and gets broader for the more stringent mapping (minAS=0.99, scores between −1.1 and 0.60). The variability of the samples for each parameter choice is high (grey violin plot **Figure 6B**), while the individual samples have much lower variance across the parameter space (purple violin plot **Figure 6B**), implying that each sample’s biologically relevant features is broadly robust across the different alignment parameters. Even when varying between the default and the most stringent values (**Figure 6C**), the co-expression scores that characterize the samples are only modestly changes, and this is, in fact, the highest level of variability observed. Generally, any pair of alignment choices yield similar co-expression scores across samples (**Figure 6D**, 0.89<r<0.99) with only the two most stringent parameter values showing any substantial deviation from the identity line. The lack of an effect on co-expression is a useful result: there is little impact on this general downstream biological application and where there is impact, it is apparently in the non-functional ranges. For most standard applications, alignment choice does not have a strong effect. It is in the functionally unannotated and novel or unknown transcriptionally active regions that researchers may need to exercise greater caution.

**Figure 6.**
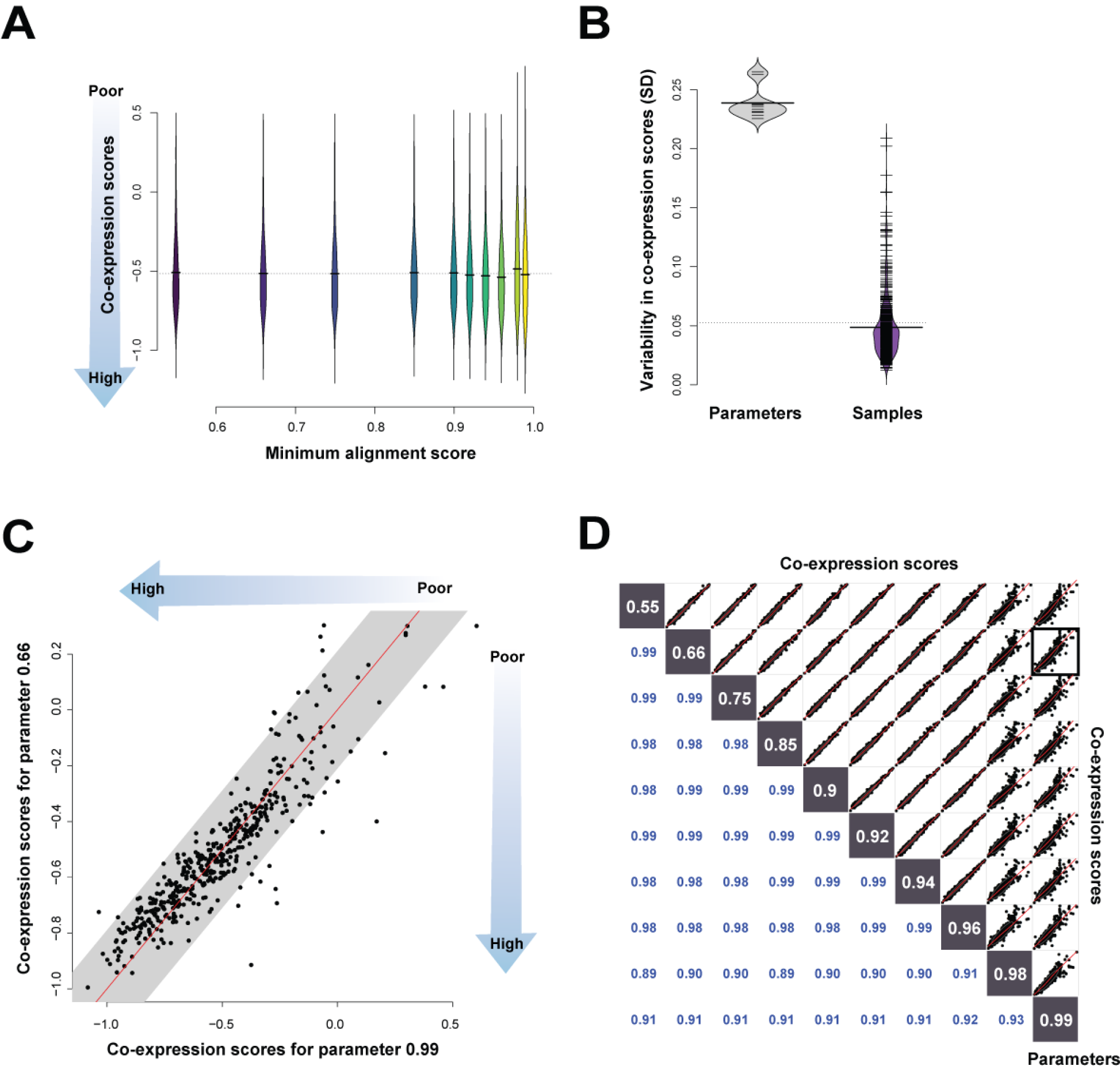
Alignment parameter impact on co-expression. (A) Distribution of co-expression scores for the different alignment parameters. (B) Differences in SDs when comparing across parameters or across samples, with more between sample variance than between parameter variance. (C) Scatterplot comparing the extreme parameter to default. Each point is the co-expression score of a sample. The grey is the average SD of the whole experiment and the identity line in black. (D) Scatterplots of each parameter against another, each point representing a sample, and each matrix square a comparison between parameter choices for the minimum alignment parameter.

Returning to the 57 experiments across the three databases, we repeat the co-expression score analysis (with the 3,405 samples, **Figure S5A-F,** SD +/-~0.32-0.33 across samples, SD +/-~0.11 across methods per sample). We see fairly similar performances across all databases on a per sample basis (correlations rs=0.73-0.95) and even higher on a per experiment basis (rs=0.92-0.95). Once more, the default STAR performances fit within expected ranges. Broadly, this seems to confirm the view that while there is substantial error and variability between pipelines, most of it does not easily alter the degree to which biological signals can be seen within the data.

### Co-expression and technical performance do not “align”

Consistent with our sex-specific DE results, on a per sample basis, we find no correlation between the co-expression replicability score and any of A) the fraction mapped (**Figure 7A**), B) the sample-sample correlations (**Figure 7B**), and C) total genes detected (**Figure 7C**). The fraction mapped metric, as previously discussed, is similar across most of the parameters, while the majority of the poorly mapped samples are for the 0.99 minAS parameter. We also find very few of the good co-expression scoring samples with very poor mapping. These trends are similar for the correlations (**Figure 7B**), since we’ve seen that the fraction mapped and correlation metrics themselves are fairly correlated. A very low positive correlation exists between the number of genes detected and the co-expression score (**Figure 7C**), whereby good co-expression scores fall within a small range of the genes detected values. Not surprisingly, this discordance between technical and biological metrics is also apparent in the corpus of RNA-seq experiments we assessed (**Figure S5D-E**).

**Figure 7.**
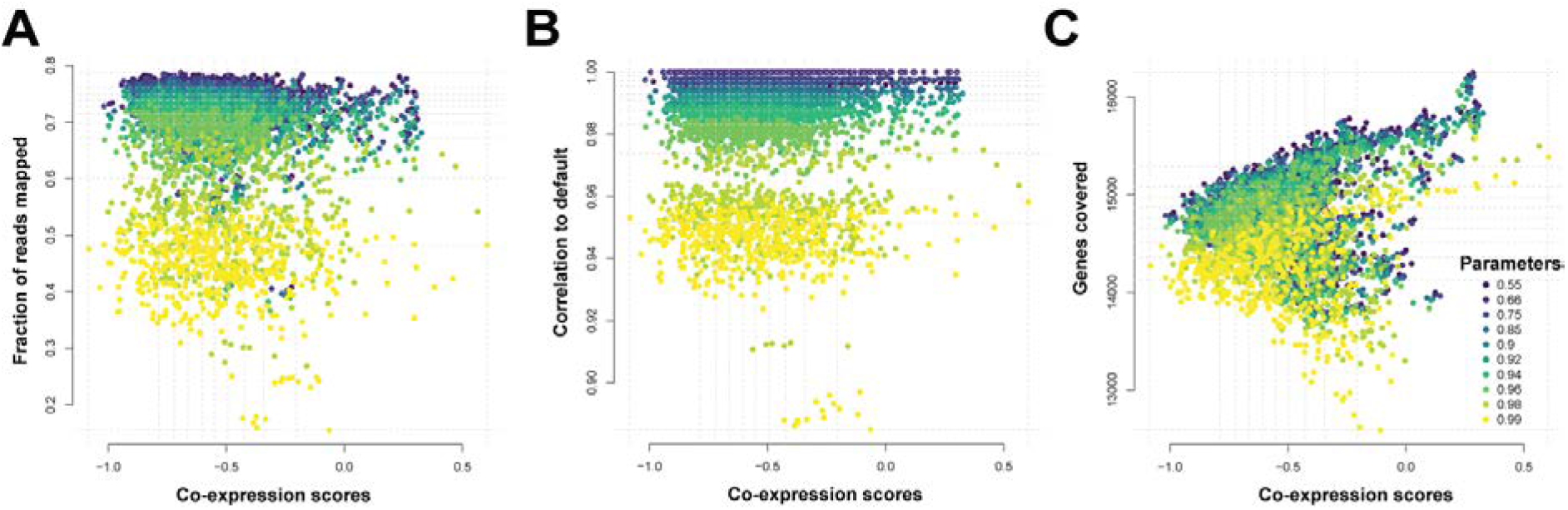
Four score and ten alignment parameters. Scatterplots comparing the co-expression score to (A) the fraction mapped (rs=-0.12), (B) correlations (rs=0.001) (C) and gene detection (rs= 0.13). Each color represents the alignment parameter tested. The grey dotted lines represent the quartiles of the scores and metrics. The co-expression scores do not correlate with the technical metrics, but are influenced by number of genes.

### Gene drop-offs caused by variation in parameters are unlikely to have a functional impact

To further explore the difference between “expressed” and “not expressed” as a function of alignment parameters, we can look at the genes that are no longer expressed (‘drop outs’) across the parameters on a per sample basis. For most genes that do lose expression or drop out, these genes were originally associated with low counts, as shown by occurrence versus average CPM values (example sample ERR188479 in **Figure S6A**). There are a few exceptions to the stricter alignment removing only low count genes observation and this was primarily in the HLA genes (e.g., *HLA-DQB2* see **Table 2**). These could potentially be related to the natural genetic variation of these regions, their splicing, or other isoform related differences (25). However, most of the functional tests and tasks we performed are not affected by these genes.

**Table 2.** Drop out genes between default and extremal parameter

| Gene | Number of Samples |
| --- | --- |
| *HLA-DQB2* | 101 |
| *HLA-DRB5* | 93 |
| *HLA-DQB1* | 43 |
| *EIF5AL1* | 36 |
| *HLA-DQA2* | 28 |
| *HLA-G* | 8 |
| *RPSAP58* | 4 |
| *PABPC3* | 2 |
| *FKBP1C* | 1 |
| *POLR2L* | 1 |
| *FRG1B* | 1 |

Although we report reads mapped uniquely, we are strictly looking at reads that map to genic or annotated regions, but there are still many reads unaccounted for. Where, then, do those reads map? If we look at the fraction of mapped reads that are mapped with the STAR “no features” label i.e., unannotated regions of the genome (comprising inter-genic and intronic regions), this number decreases slightly as we increase the stringency (**Figure S7A**). So, although we are mapping more to the genome under the more permissive parameters, we are principally mapping to less functionally annotated and also non-exonic regions, and therefore it should not be surprising that our assessments that rely on functionally annotated genes are affected. Although these fractions are fairly small (averaging near 5%), these reads are not contributing to any gene or function that can be assessed in an experimentally useful fashion (see supplementary text for additional discussion).

### Filtering of genes post-alignment does alter performance

Although we find that parameter changes which toggle a small number of genes with low counts to zero (or back) have no major effect on performance, we know from the MAQC/SEQC benchmarking that filtering away low expressing genes generally improves replicability (26). To better explain this apparent discrepancy, we repeat our analysis holding all parameters constant at default but now filtering away low expressing genes at different count thresholds. Further to this, we repeat the analysis varying the mismatch parameter (nM) and downsampling total reads to simulate lower depth experiments. These three features represent important choices in sequencing and alignment: the recommended post-alignment filter, effect of sequencing errors and potential variation (SNPs, indels or SVs) and the effect of read depth. For the mismatch parameter, as with the minimum alignment score parameter, the more stringent the parameter, the less reads mapped but the gene detection does not vary as highly. Even though reads mapped ranged on average between ~13M and 22M reads (min 3M, max 63M), gene detection only changed between 14.2K and 14.9K (min 12.5K, max 16.2K **Figure S8A-C**). Not surprisingly, filtering away genes with low counts changed gene detection the most (**Figure S8D**). This filtering is far larger in degree than the modest changes induced indirectly by changing most parameters, explaining its greater impact. Alignment is robust well outside the range of expression values that are plausibly biologically important, requiring an independent basis for filtering.

We then calculate the co-expression scores across the above parameter choices and plot the distributions. As before, we see similar scores across the parameters (only extreme parameters shown **Figure 8A,** all parameters in **Figure S9**) with the more stringent parameters almost identical, except when filtering away low expressing genes as reflected in the larger spread in the distribution of scores. This affirms the “low expressing genes are mostly noise” argument. Because downsampling reads does not have this effect (first violin plot pair in **Figure 8A**), this indicates that lower sequenced read depth, if randomly distributed, is less of a problem compared to low expressing genes. We then tested the effect of both alignment and filtering by calculating co-expression scores across both parameters, averaging the runs across samples and interpolating between points (**Figure 8B**). For the most part, the landscape is flat, with most parameters performing equally, with some variation (e.g., stringent minAS, extreme filtering). This is also consistent in a second dataset (**Figure S10**).

**Figure 8.**
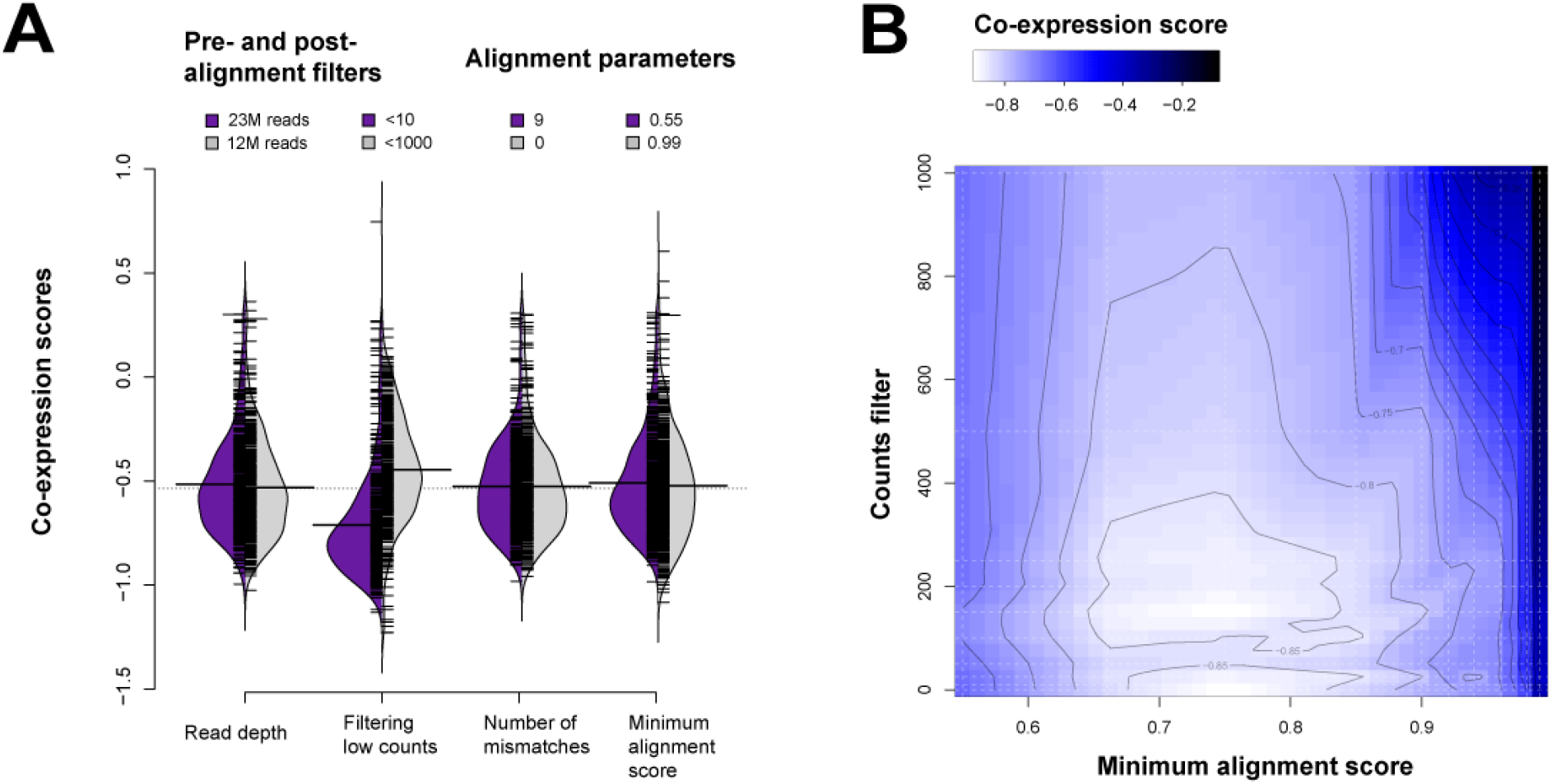
Defining the parameter landscape of STAR. (A) Distributions of scores for the extreme parameters tested on the GEUVADIS dataset. The purple distribution shows the scores per sample of the most permissive parameter and the grey distribution the most stringent. Downsampling reads has little effect on the co-expression scores. Filtering low expressing genes by counts has the greatest impact, consistent with the SEQC replicability experiments. Allowing for mismatches has little impact. The minimum alignment score parameters are also placed for comparison. (B) Interpolated co-expression scores showing alignment and post-alignment filter hotspots (light blue to white). Dark areas are least replicable. These results are averaged over all 462 samples for 1000 runs, and interpolated between the dashed lines. Contours define interpolated score boundaries.

## DISCUSSION

Our results paint a picture of an interesting landscape: in technical respects, alignment works well and similarly across algorithms, until it dramatically fails, with the point of transition being quite extreme in most data. In addition, much of the technical variation in performance has little direct bearing on the degree to which biology can be intuited from the results. We like the analogy of ice floes for this scenario, in which the landscape is relatively flat across a wide range of parameters, until one falls off the surface, with a hard-to-assess ground landscape always below (as suggested by **Figure 8B**). This view is consistent with our observation both of the variation in performance across algorithms in many individual assessments and datasets, where the most notable variation in performance was in occasional outliers, and also with our in-depth testing-to-destruction of STAR. While the failure modes observed in STAR are not certain to generalize across methods or other parameter choices we have not explored, the similarity in results obtained in our cross-database analysis do suggest broad applicability for our findings. In essence, we find that gene expression is robust except where it collides with a requirement to know precise sequence (e.g., X-Y paralogs) or obtain exact values (e.g., expressed versus not) rather than statistical differences or trends. These are both the potential fractures within alignment performance and, not coincidentally, constitute the major avenues of ongoing research (e.g., allele-specific expression, single-cell drop-out, novel isoforms, etc).

However, for estimating gene expression, and in a purely technical sense, alignment assessment is subject to Fredkin’s paradox, in which the difficulty of picking a winner reflects the broad similarity in output. Where this technical capacity is biologically insufficient (e.g., subtle changes in gene expression do matter) is not trivial to say. Moving past technical performance to an assessment of biology will require a paradigm shift from developers, more narrowly targeting what users are interested in. The best current assessments rely on either simulated datasets with a known ground truth (e.g., compcodeR (27), or (20)), real datasets with replicates (e.g., rnaseqcomp (6), URL: http://rafalab.rc.fas.harvard.edu/rnaseqbenchmark), or real datasets with complementary expression values (e.g., nanostring results as in RNAontheBench (28), or qPCR results as with the MAQC data (26)). Unfortunately, even using real expression values does not trivially solve the problem of assessment with gold standards that can generalize across datasets being hard to come by. Sex-specific analyses offer a clear way forward, being both general (applicable to many datasets) and biological; its drawback, however, is that this particular test is too easy. And, of course, our own assessment does not encompass all the use-cases of RNA-seq, particularly those that require knowing absolute expression levels or precise sequences. Establishing a few harder ground truth assessments from biological data should be possible, exploiting, for example, tissue or cell-type specificity. While purely algorithmic developments should be encouraged, it is critical to distinguish them from improved biological assessment, which could be validated with a novel observation within pre-existing data, particularly meta-analytically.

It shouldn’t be a surprise, perhaps, that most tools perform reasonably well with respect to simple metrics: they are designed to. What, then, should assessments and developers focus on? We think understanding the factors leading to differences in performance is far more important than obtaining good performance. All developers conduct internal validations which optimize the performance of their own tool; reporting all these performances has more utility than is currently appreciated. While characterizing a range of performances diminishes the degree to which new algorithms can be said to “beat” others, we suspect it more accurately reflects that methods work by being optimized for different tasks, which sometimes involves quite real trade-offs. Knowing those trade-offs and making informed decisions would be useful. Our second suggestion is for a greater appreciation that many technical metrics may show clear signals precisely in the domain where there is little variation in biological signal, whether because it is too strong (sex DE) or too weak (low expression). In addition to these potential targets for improved output, improvements in process alone through computing performance (i.e., memory usage, speed) are both easier to quantify and highly desirable.

Throughout this work, we have focused on assessments and assessors, motivated by an interest in better understanding variation (or lack thereof) in RNA-seq alignment. However, one question that still remains is what should users do when faced with true alignment variation across tools? Or any bioinformatics tool for that matter? As in most software development, a report of such cases to developers would be ideal. If developers maintain a public record of such cases, and enough are collected with similar issues, it then becomes a community problem to be solved. And as in the case in competitions for gene function prediction (e.g., CAFA (29)) or more general systems biology questions (e.g., DREAM challenges (30)), large-scale efforts will bring to light the issues and garner creative solutions. More broadly, thoughtful reporting and characterization of edge cases will provide both users and developers with information to understand current performance and optimize pipelines for specific biological interests where alignment remains challenging.

## AVAILABILITY

AuPairWise is available in the GitHub repository (https://github.com/gillislab/aupairwise)

STAR is available in the GitHub repository (https://github.com/alexdobin/STAR/).

## SUPPLEMENTARY DATA

Supplementary Data are available at NAR online.

Additional code is available in the GitHub repository (https://github.com/gillislab/aligner)

